# Neuropeptide NLP-40 and Cognate G-Protein Coupled Receptor AEX-2 Regulate Behavioral Responses During Anoxia in *Caenorhabditis elegans*

**DOI:** 10.64898/2026.09.30.755124

**Authors:** Ginger Z. Watzinger, Heather L. Bennett

## Abstract

**Background:** Insufficient oxygen (hypoxia) or complete lack of oxygen (anoxia) causes cellular dysfunction and death in terrestrial animals. As a result, animals have developed homeostatic strategies, including changes in movement strategies to escape and survive stressful environments. *Caenorhabditis elegans* respond to changes in oxygen by increasing the frequency of reversals (backward movement) and increasing locomotor speed; these behaviors enable them to escape unfavorable environments and move to more suitable ones. The molecular and neural mechanisms underlying behavioral responses to low-oxygen environments are incomplete. Neuropeptides regulate *C. elegans* behaviors by modifying neural circuit function in response to hypoxia. Loss of function in *C. elegans* neuropeptide *nlp-40* and its receptor *aex-2* increased survival to anoxia [1]. However, the cells in which NLP-40 and AEX-2 function for anoxia, as well as the mechanisms underlying responses to anoxia, remain elusive. We hypothesized that NLP-40 and AEX-2 regulate anoxia by altering escape behaviors.

**Results:** We found that neuropeptide NLP-40 functions in cholinergic neurons and AEX-2 functions in cholinergic or GLR-1 interneurons or motor neurons to regulate susceptibility to anoxia. We found that loss of either *nlp-40* or *aex-2* decreased locomotor activity and reversal frequency early during 48 hours of anoxia, suggesting that *nlp-40* and *aex-2* are required for the proper execution of these behaviors. We found that neuronal knockdown of *nlp-40* recapitulated the behaviors observed in *nlp-40* mutant animals and increased survival; however, restoring *aex-2* function in *glr-1* interneurons or motor neurons was sufficient to rescue behaviors to wild-type and restore vulnerability to anoxia.

**Conclusions:** Overall, our results suggest that NLP-40 and AEX-2 likely function in cholinergic or *glr-1* interneurons or motor neurons to regulate escape behaviors during anoxia. Escape behaviors are part of the homeostatic response and are typically protective in the wild; however, in our paradigm, we show that loss of either *nlp-40* or *aex-2* reduces escape responses and confers protection against 48 hours of anoxia.

Collectively, these results highlight a new role for NLP-40 and AEX-2 in anoxia and shed light on mechanisms underlying the adaptive responses to anoxia.

## Introduction

Oxygen is the final electron acceptor in cellular respiration and is therefore essential for sustaining all biological processes in most metazoans [2–4]. Hypoxia or anoxia challenges animals, as they must adapt to this stress or die. As a result, animals have developed multifaceted, complex strategies to maintain cellular homeostasis that alter physiology, neural activity, and behavior [5–7]. Animals generate different behaviors and use different movement strategies depending on internal cellular conditions, environmental cues, and established behavioral experiences [8–12]. Substantial progress has been made in defining the cellular and molecular mechanisms that detect changes in oxygen availability [13–15]. Less is known about the molecular and neural mechanisms that coordinate behavioral changes in response to low-oxygen conditions, an important and open area of research [16].

The nematode *Caenorhabditis elegans* has been an invaluable model for interrogating the molecular and neural mechanisms underlying behavioral responses to low-oxygen conditions across diverse tissues and different developmental stages [15]. In response to shifts in oxygen concentrations, *C. elegans* exhibit behavioral changes marked by a brief increase in forward locomotion, increased reversal and turning frequency, and a subsequent reduction in overall activity [12, 17–19]. In the wild, oxygen is unevenly distributed in soil environments, and *C. elegans* must monitor internal and external oxygen concentrations to maintain cellular homeostasis [20], suggesting these behaviors may be protective, allowing *C. elegans* to navigate changes in oxygen gradients and escape to find more favorable conditions. *C.elegans* prefer oxygen environments between 5 and 12% and possess oxygen-sensing neurons that detect changes in environmental oxygen and engage neural circuits to regulate escape behaviors [17–19, 21–24]. These behavioral changes can be observed in hyperoxia (>12% oxygen) or with different hypoxic states (0.5-1% oxygen available) [25, 26]. Studies of oxygen-dependent behavior in *C. elegans* have shown that acute and prolonged hypoxia can modify neural circuit function, and neuromodulators, such as neuropeptide signaling, contribute to these responses [17, 18, 27–30]. Specific neuropeptide signaling pathways have been implicated in driving hypoxia-dependent behavioral responses, modifying escape behaviors such as locomotion and reversal turns [28, 30]. During shifts in oxygen availability, specific interneurons release neuropeptides to regulate these locomotory responses and signal to sensory neurons. [31]. Beyond their role in modulating hypoxia-dependent behaviors, under normoxia, with ample oxygen available, mutations in neuropeptide signaling perturb locomotion and overall movement, further emphasizing neuropeptides in mediating behavioral responses [32, 33]. Collectively, these studies and others support a role for neuropeptide signaling in mediating behavioral adaptation to sustained environmental challenges such as low oxygen conditions.

Furthermore, neuropeptide signaling mediates sensitivity and resistance to diverse hypoxic states. *C. elegans* deficient in the neuropeptide proprotein convertase gene *egl-3* and the carboxypeptidase E gene *egl-21,* as well as the gene *unc-31,* which encodes a calcium-dependent activator protein, were resistant to anoxia. [34]. A follow-up study showed that nervous system-specific rescue or knockdown of *egl-3* restored susceptibility or increased resistance to anoxia, respectively [1]. Moreover, loss of neuropeptide *nlp-40* and its putative cognate G protein-coupled receptor *aex-2* mimicked loss of *unc-31* and *egl-3* and also increased survival to 48 hours of anoxia. Like *egl-3*, inhibition of *nlp-40* gene function broadly in neurons increases survival to anoxia, suggesting that NLP-40 also functions in neurons to regulate responses to anoxia [1]. The mechanism(s) by which neuropeptide *nlp-40* and the *aex-2* receptor regulate anoxia and confer protection remains to be determined. Moreover, it is unknown whether *nlp-40* and *aex-2* function, like other neuropeptides, to coordinate behavioral responses to low oxygen conditions.

Here we investigate the cellular and molecular mechanisms by which the neuropeptide *nlp-40* and its putative cognate G protein-coupled receptor *aex-2* regulate behavioral responses to anoxia. We used classic genetic manipulation approaches of *nlp-40* and *aex-2* with behavioral analyses and found that loss of either *nlp-40* or *aex-2* decreased overall locomotor activity and reduced reversals. This conferred protection and increased survival during anoxia. We found that NLP-40 signals from cholinergic neurons and AEX-2 likely signals from interneurons or motor neurons render animals susceptible to anoxia, and restoring gene function is sufficient to restore behavioral responses, with changes in body bends and reversals. Our results highlight a new role for neuropeptide *nlp-40* and the *aex-2* receptor in mediating escape behaviors in response to anoxia.

## Materials and Methods

### *C. elegans* maintenance and media preparation

Strains were reared and assayed at 20 degrees Celsius, unless otherwise stated, on nematode growth medium (NGM plates) seeded with *Escherichia coli* OP50 as a food source under standard conditions [35]. NGM plates were seeded with 40 µL of OP50. Six-well plates for body bends and reversal assays were prepared with 5mL of NGM and seeded with 10 uL of bacteria the night before the assays.

### Strains

The following strains were used in this work:

N2, referred to as wild type

VC671 *egl-3(n979)* V

CB928 *unc-31(e928) IV*

NM4394 *nlp-40(tm4085)*, 0x backcrossed

OJ1188 *nlp-40(vj3) IV*

HLB14 *nlp-40(tm4085) IV*, 2x backcrossed

JT3 *aex-2(sa3) X*

PS7837 *aex-2(sy1078) X*

NM3242 *aex-2(sa3)X*;jsEX937[*aex-2p::aex-2::mCherry*]; *pPD118.33[Pmyo-2::GFP]]*

NM3228 *aex-2(sa3)X;jsEx925[(Pmyo-3::aex-2::YFP); pPD118.33[Pmyo-2::GFP)]*

NM3232 *aex-2(sa3)X;jsEx929[(Pges-1::aex-2::GFP);pPD118.33(Pmyo-2::GFP)]*

NM3389 *aex-2(sa3)X;jsEx999[(Pglr-1::aex-2::GFP); pPD118.33(Pmyo-2::GFP)]*

NM3397 *aex-2(sa3)X;jsEx1007[Punc-17::aex-2::GFP);pPD118.33(Pmyo-2::GFP)]*

NM3393 *aex-2(sa3)X;jsEx1003[(Punc-47::aex-2::GFP);PD118.33(Pmyo-2::GFP)]*

*TU3311 uIs60 [unc-119p:: YFP + unc-119::sid-1]*

*XE1581 wpSi10 II; eri-1(mg366) IV; rde-1(ne219) V; lin-15(n744) X*

*XE1582 wpSi11 II; eri-1(mg366) IV; rde-1(ne219) V; lin-15(n744) X*

### Anoxia Survival Assay using AnaeroGen Gas Generator System

Gravid adults were synchronized using a bleach hypochlorite treatment as described in Stinergnale *et al.,* and used in anoxia and hypoxia assays in previous studies [36, 37]. Anoxia experiments were done with L4-stage animals. For anoxia exposure, 30-40 early-stage and mid-L4-stage animals per genotype were selected and exposed to 48 hours of anoxia using an Oxoid AnaeroGen W-Zip Compact Gas Generator System^TM^ (anaerobic environmental system, with an Oxoid Resazurin Anaerobic Indicator^TM^ (Thermo Scientific). Animals were exposed to 48 hours of anoxia at 20°C, then re-exposed to oxygen at 20°C for 24 hours. After 24 hours, animals were scored for survival. Survival was determined for animals that developed into adults, regained movement, and/or resumed feeding 24 hours after re-oxygenation as previously described in Doshi et al., 2019.

### RNAi Experiments

NGM plates were prepared with either *E. coli* expressing the *L4440* empty vector or *nlp-40 RNAi* encoding Y74C9A.2 plasmids for feeding RNAi experiments, purchased from Horizon Discovery. L4 stage animals were placed on either L4440 or *nlp-40* RNAi plates for at least four generations. Gravid adults were bleach-prepared to synchronize animals. Fourth-generation L4 stage animals were selected and placed in anaerobic environmental bags for 54 hours and then exposed to oxygen for 24 hours before scoring survival or behavior. RNAi neuron-sensitive animals were reared and assayed at 20 degrees Celsius, except XE1581 animals, which were maintained at 15 degrees and tested at 20 degrees Celsius.

### Scoring body bends and reversal turning frequency

Body bends were scored under room air, normoxic conditions. 10 larval (L4) stage animals of each genotype were picked to NGM agar seeded with *OP50* bacteria the day before. For body bend experiments conducted under normoxia, animals were allowed to recover and acclimate to room air conditions for three minutes before scoring for body bends. The total number of body bends is scored continuously for 3 minutes. The average number of body bends per minute for each worm is calculated. For body bend and reversals experiments conducted under anoxic conditions, using the AnaeroGen W-Zip Compact Gas Generator System, (L4) stage animals are tested in six-well plates seeded with *E.coli OP50* bacteria the day before. Three animals per well are selected to a well, for a total of nine animals for each genotype. Age- and stage-matched animals are tested as controls. Animals are placed into the six -well assay plates and allowed to acclimatize to laboratory and AnaeroGen bag environments for 10 minutes. Body bends and reversal turns were subsequently scored in the same assay at 30 minutes and 60 minutes after the start of anoxia. For behavioral tests using RNAi-fed strains, animals were reared and maintained as described above. NGM media for 6-well plates were prepared using standard RNAi protocols as described in [38]. Wells were seeded with either empty vector bacteria or n*lp-40 RNAi*, the day before the assay. For most behavior experiments or trials, samples were numbered randomly so that the genotype was blinded when data were collected; the samples were not decoded until after all data were collected.

### Statistical Analysis

Statistical analysis was performed using GraphPad Prism version 11 (GraphPad Software, La Jolla, CA). For anoxia experiments done using the AnaeroGen W-Zip Compact Gas Generator System, these experiments were done in triplicate with 30 to 40 animals per genotype, and independent trials were done on different days; as a result, survival results were pooled from data collected from at least three independent trials. Statistical significance was determined using a one-way ANOVA followed by a Tukey’s or Bonferroni multiple comparisons post hoc test. For comparisons with two genotypes, we used a t-test. We report the standard error of the mean for all experiments. Body bends and reversals assessing basal activity and turning behaviors were scored for each genotype and condition with at least three replicates, with experiments conducted on independent days. Statistical analysis for body bends was assessed using a two-way ANOVA with a Sidak correction. Significance was considered if p< 0.05. All data, including the number of animals scored and the number of independent replicates, are available in the supplement or upon request.

## Results

### Loss of Either *nlp-40* or *aex-2* Increases Survival to Anoxia

We were interested in elucidating the role of neuropeptide *nlp-40* and cognate receptor *aex-2* in regulating behavioral responses to anoxia. A previous study showed that loss of function in either neuropeptide *nlp-40* or its cognate G protein-coupled receptor *aex-2* confers protection and increases survival to anoxia [1]. First, we aimed to independently reproduce previously reported findings by testing two different loss-of-function alleles in neuropeptide *nlp-40*: *nlp-40(tm4085)* and *nlp-40(vj3)*; both mutations delete large regions within the *nlp-40* coding sequences and are null alleles [39]. We recapitulated published results: loss in neuropeptide NLP-40 leads to an increase in survival to 48 hours of anoxia (Figure 1A-C; N2 0.44 +/-0.05 versus *nlp-40(tm4085)* 0.68 +/-0.04, ANOVA *F= 14.88,* p value 0.004; N2 0.46 +/-0.04 versus *nlp-40(tm4085) 2x backcross* 0.73 +/-0.02, ANOVA *F= 33.89,* p value < 0.0001; N2 0.58 +/-0.04 versus *nlp-40(vj3)* 0.82 +/-0.03, *F= 19.35,* p value < 0.0001). Next, we tested *aex-2*, a G protein-coupled receptor gene and cognate receptor for *nlp-40*. Consistent with previous findings from Doshi et al., we found that *aex-2(sa3),* a strong loss-of-function allele, resulted in increased survival to 48 hours of anoxia (Figure 1D; N2 0.55 +/-0.04 versus *aex-2(sa3)* 0.68 +/-0.03, ANOVA F() = 36.40, p value 0.03). We extended our analysis and tested an additional allele of *aex-2*, *aex-2(sy1078)*, a CRISPR-generated null allele [40], and found *aex-2(sy1078)* animals also showed increased survival and conferred protection to 48 hours of anoxia (Figure 1E; N2 0.40 +/-0.05 versus *aex-2(sy1078)* 0.60 +/-0.05, ANOVA F() = 40.77, p value 0.04). Together, neuropeptide *nlp-40* partially regulates survival to anoxia, likely in part through its putative receptor, *aex-2.* Additionally, we have recapitulated published findings, demonstrating that our assays can detect changes in resistance and vulnerability to anoxia.

**Figure 1:**
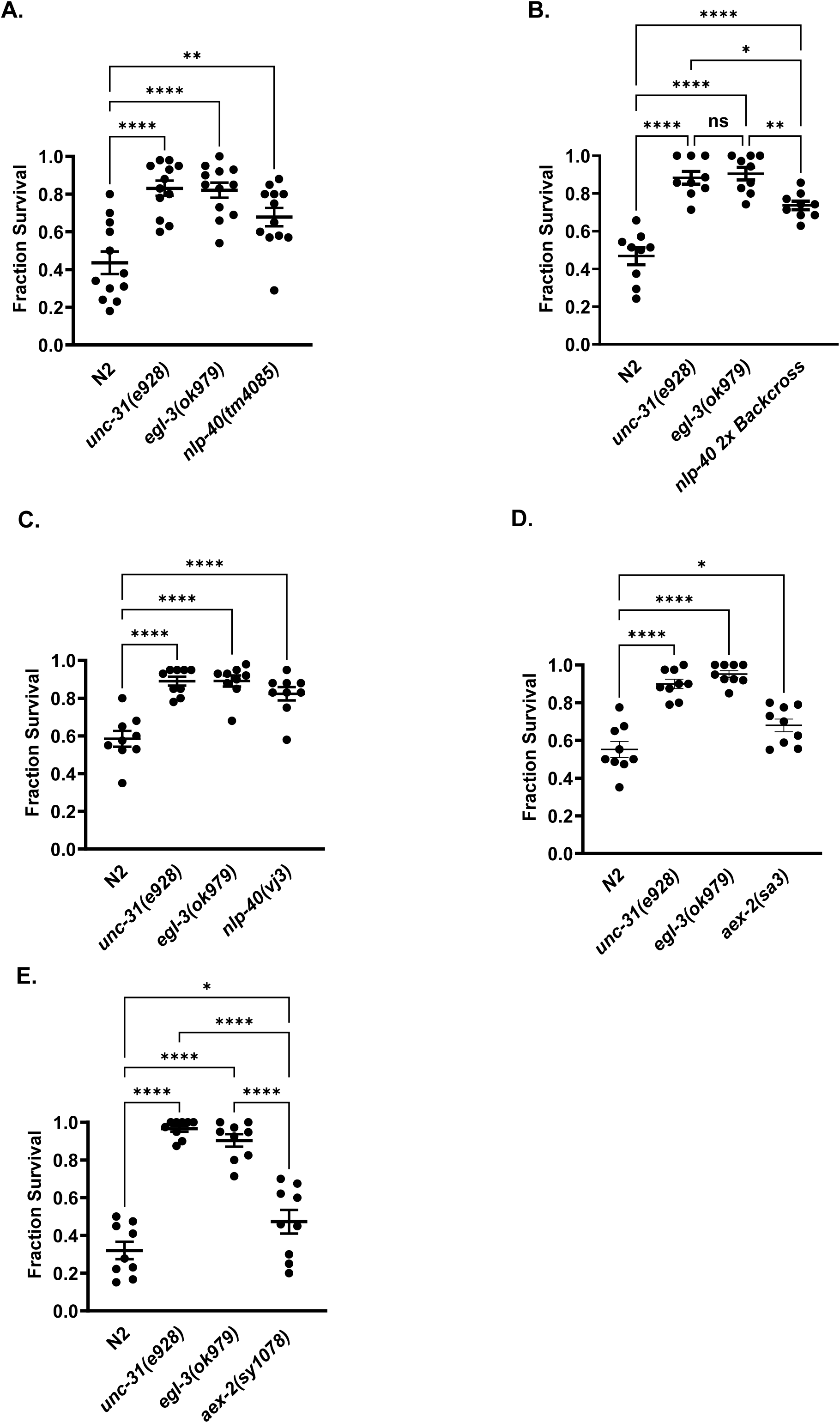
Loss of *nlp-40* or *aex-2* leads to increased survival of anoxia. **A-C) Loss of *nlp-40* increases survival to 48 hours of anoxia.** Survival of L4 stage *nlp-40(tm4085)* 0x backcrossed (b), *nlp-40(tm4085)* 2x backcrossed (c), and *nlp-40(vj3)* loss-of-function mutant animals after 48 hours of anoxia and 24 hours of re-oxygenation, compared to N2 wild-type animals. Loss-of-function in neuropeptide processing genes, *egl-3(ok979)* and *unc-31(e928),* were co-bagged and assayed as positive controls. Mean fraction of survival is reported, standard error of the mean. Results are shown for at least 4 independent trials, 360 animals per genotype. **** denotes significance with a p value <0.001, ** p value 0.004, * p value 0.02 assessed by one-way ANOVA, followed by a Tukey post hoc test. **D-E) Loss of cognate aex-2 receptor increases survival and protection against anoxia.** Survival of L4 stage (d) *aex-2(sa3)* and (e) *aex-2(sy1078)* loss-of-function mutant animals after 48 hours of asphyxiation and 24 hours of re-oxygenation compared to wild type, *egl-3(ok979),* and *unc-31(e928).* Results are shown for 4 independent trials, 360 animals per genotype. **** denotes significance with a p-value <0.0001 compared to N2 wild-type; * p value 0.03 for *aex-2(sa3)* and p value 0.04 for *aex-2(sy1078)* assessed by one-way ANOVA, with a Tukey post hoc test.

### AEX-2 Functions in Cholinergic and *glr-1* Interneurons to Regulate Anoxia

We tested animals expressing *aex-2*cDNA behind its native promoter in *aex-2(sa3)* mutant animals. L4-stage and age-matched N2 wild type, as well as *egl-3(ok979)*, *unc-31(e28),* and *aex-2(sa3)* loss-of-function mutant animals were simultaneously tested as controls. We found that restoring *aex-2* function in *aex-2(sa3)* mutant animals suppressed the increased survival observed in *aex-2(sa3)* loss-of-function animals (*aex-2(sa3*) 0.64 +/-0.07 versus *aex-2(sa3);aex-2p::aex-2* 0.37 +/-0.04, ANOVA F(4,55)=21.63, p value 0.004 (Figure 2A). This result demonstrates that loss of *aex-2* yields a protective effect and increases survival to anoxia.

**Figure 2:**
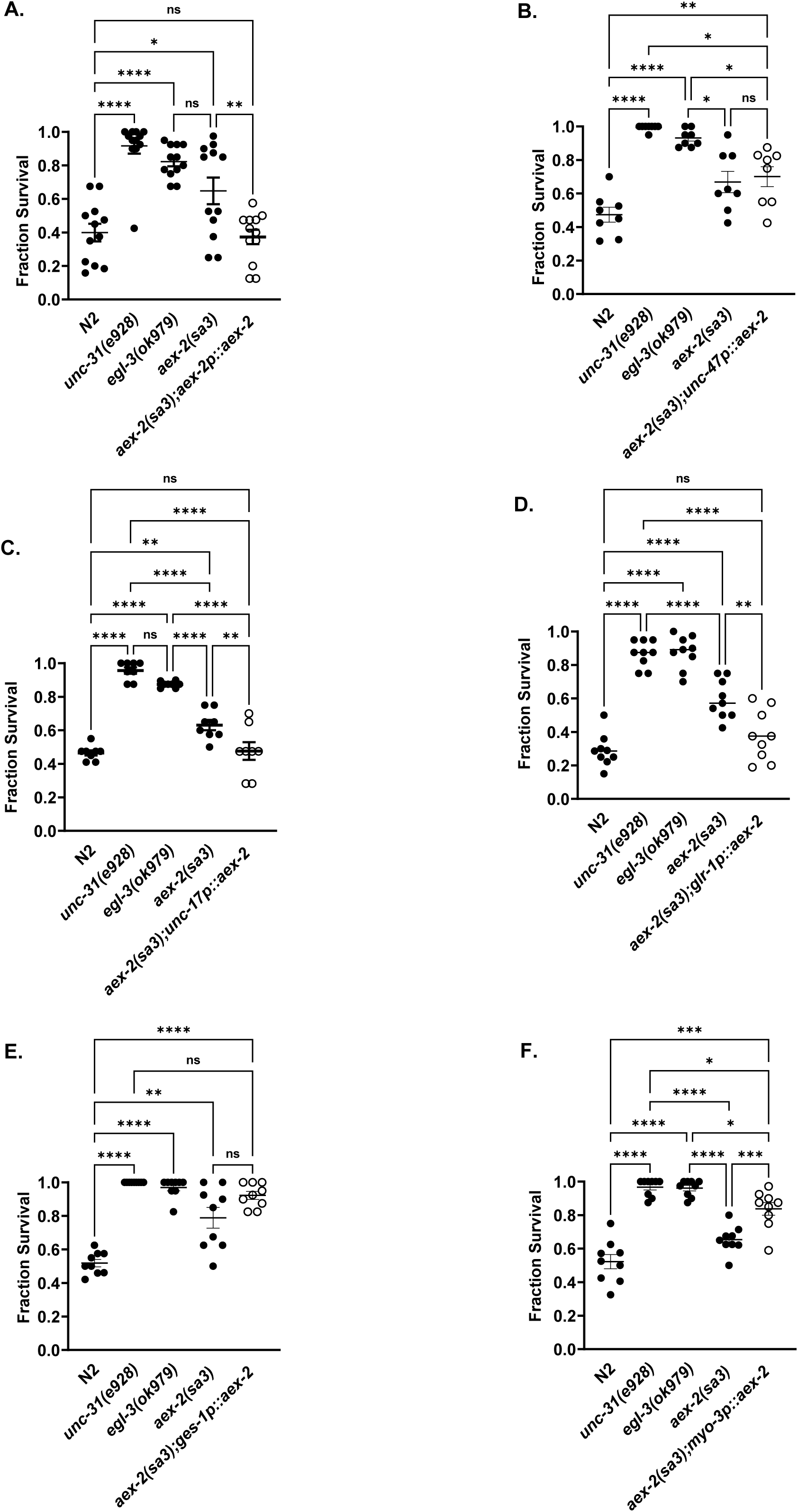
Restoring *aex-2* gene function in neurons is sufficient to restore vulnerability to anoxia. **A) Restoring *aex-2* gene function under the *aex-2* endogenous promoter restores sensitivity to 48 hours of anoxia.** Survival of L4-stage *aex-2(sa3)* animals expressing rescue copy of *aex-2cDNA* behind the endogenous *aex-2* promoter (open circles) exposed to 48 hours of anoxia, followed by 24 hours of re-oxygenation. Mean fraction of survival is reported and compared to wild type, *aex-2(sa3)*, *egl-3(ok979),* and *unc-31(e928); (lf) mutant* animals (shaded circles). Standard error of the mean is recorded. Results are shown for 4 independent trials, n=360 animals per condition. Statistical significance assessed by one-way ANOVA, **** p value <0.0001, ** 0.05, * 0.01 Tukey post hoc test. **B) AEX-2 does not function in GABAergic neurons or non-neuronal tissues to regulate response to anoxia.** *aex-2(lf)* animals expressing aex-2cDNA behind the GABAergic promoter *unc-47* tested as in panel A. Results are shown for 4 independent trials, 360 animals per genotype. * denotes statistical significance, **** p value <0.0001, ** 0.008, * 0.01. **C) Restoring AEX-2 Function in Cholinergic Neurons Increases Susceptibility to Anoxia.** *aex-2(lf)* animals expressing rescue copy of *aex-2cDNA* behind the *unc-17* promoter tested as in panel A. Results are shown for 4 independent trials. * denotes statistical significance, **** p value < 0.001, ** 0.003 N2 compared to *aex-2(sa3)* and ** p value 0.006 *aex-2(sa3)* compared to *aex-2(sa3);unc-17p::aex-2*. **D)** AEX-2 Functions in *glr-1* Expressing Neurons to Mediate Response to Anoxia. Survival of L4-stage *aex-2(lf)* animals expressing a rescue copy of *aex-2 cDNA* under the *glr-1* promoter was tested and scored for survival as in panel A. Results are shown for 4 independent trials. * denotes statistical significance assessed by one-way ANOVA, * denotes statistical significance, **** p value <0.0001, ** 0.001. **E-F) AEX-2 Is Not Required in Non-neuronal Tissues to Regulate Anoxia.** *aex-2(sa3lf)* animals expressing aex-2cDNA behind (e) body wall promoter *myo-3*, or (f), an intestinal promoter *ges-1* tested as in panel A. Results are shown for 4 independent trials, 360 animals per genotype. * denotes statistical significance, **** p value <0.0001, ** p value 0.009 N2 compared to aex-2(sa3), *** p value 0.004, * p value 0.03.

Next, we asked which tissue(s) *aex-2* functions to regulate anoxia. We hypothesized that *aex-2* likely function in neurons to confer protection against anoxia. This postulation is based on the following findings: (1) Neuropeptide processing genes *egl-3*, *egl-21*, and *unc-31* are ubiquitously expressed in the nervous system [41]. Restoring or reducing *egl-3* function throughout the entire nervous system in L4-stage animals suppressed or increased survival to anoxia, respectively [1]. (2) Fluorescent reporter analysis indicates that AEX-2 is expressed in neurons [42]. (3) AEX-2 functions in two GABAergic neurons, AVL and DVB, to regulate defecation, bolstering support for a neuronal role for AEX-2 in regulating *C. elegans* behavior [39]. As a result, we tested *aex-2(sa3)* mutant animals expressing an *aex-2cDNA*::GFP fusion behind *unc-47* GABAergic promoter. However, we found that restoring *aex-2* function in GABAergic neurons failed to suppress the increase in survival to 48 hours of anoxia (*aex-2(sa3*) 0.69 +/-0.06 versus *aex-2(sa3)unc-47p::aex-2* 0.70 +/-0.05; ANOVA F(7,28)=30.37, p-value 0.99. (Figure 2B). This result suggests that the neurons and genetic pathways where *aex-2* functions for defecation are likely distinct from those involved in anoxia.

Because fluorescent reporter analyses also detect *aex-2* expression in cholinergic and *glr-1* interneurons [42], we tested if animals expressing restored *aex-2* function (*aex-2*cDNA GFP fusion) behind the *unc-17* promoter or *glr-1* promoters, which drive expression in cholinergic or a subset of interneurons, would suppress increased survival to anoxia. We find *aex-2* function in either cholinergic or *glr-1* interneurons suppresses survival and increases susceptibility to anoxia, similar to wild type (*aex-2(sa3*) 0.63 +/-0.03 versus *aex-2(sa3);unc-17p::aex-2* 0.47 +/-0.05, ANOVA F(4,35)=58.03, p value 0.006), (*aex-2(sa3*) 0.59 +/-0.03 versus *aex-2(sa3);glr-1p::aex-2* 0.37+/-0.05, ANOVA F(4,40)=52.65, p value 0.001) (Figures 2C-D).

To exclude the possibility that *aex-2* signals from non-neuronal tissues also contribute to the response to anoxia, we tested animals that express *aex-2*cDNA GFP in either the intestine or in body wall muscles, using *ges-1* and *myo-3* promoters, which drive expression in the intestine and body wall muscles, respectively. However, we find neither *aex-2* expression in the intestine or body wall muscles failed to suppress the increase in survival to anoxia (*aex-2(sa3*) 0.79 +/-0.01 versus *aex-2(sa3);ges-1p::aex-2* 0.92 +/-0.06, ANOVA F()=36.99, p value 0.11) (*aex-2(sa3*) 0.65 +/-0.02 versus *aex-2(sa3);myo-3p::aex-2* 0.83 +/-0.03, ANOVA F () = 3.804, p value 0.0007 (Figure 2 E-F).

Collectively, our results support a model in which *aex-2* functions in a non-cell-autonomous manner, within a subset of cholinergic or *glr-1* interneurons, to regulate response to anoxia.

### NLP-40 Likely Functions in Neurons To Mediate Response to Anoxia

We asked if NLP-40 functions in specific tissues to regulate anoxia. We began testing neurons, because a previous study showed that diminished function broadly throughout the nervous system showed increased survival to anoxia that mimicked the response observed with *nlp-40* loss-of-function alleles [1]. Because neuropeptides can be readily overexpressed, and mis-expressing neuropeptides in cells where they do not function increases the likelihood of false-positive rescue or off-target effects, potentially leading to inaccurate conclusions. We decided to knock down *nlp-40* gene expression using RNA interference (RNAi). We used a *C. elegans* strain TU311, which expresses the dsRNA *sid-1* channel behind the *unc-119* pan-neuronal promoter, which enhances dsRNA uptake in the nervous system [43]. Animals were fed either control empty vector bacteria or *nlp-40* RNAi to induce knockdown of *nlp-40* in the entire nervous system. *C. elegans* expressing *nlp-40* RNAi pan-neuronal compared to those raised on empty vector showed increased survival to anoxia (Figure 3A, *sid-1* empty 0.43 +/-0.38 versus *sid-1 nlp-40* RNAi 0.80 +/-0.06, t()=5.041 p=0.001, paired t test). Our findings are consistent with previous observations that loss of *nlp-40* in neurons confers protection and increases survival to anoxia. To ascertain which neuronal populations NLP-40 functions to regulate anoxia, we used *nlp-40* RNAi in specific neuronal subtypes. Single-cell RNA transcript analysis shows *nlp-40* expression in the nervous system, with significant enrichment in cholinergic neurons [44, 45]. As a result, we prioritized testing of cholinergic neurons. Like Doshi *et al.,* we used XE1581, a *C. elegans* strain that allows for enhanced dsRNA specifically in cholinergic neurons [46]. Initially, we set out to recapitulate the results reported by Doshi *et al.,* in part to confirm our experimental design and setup worked similarly in our hands. Doshi and colleagues showed knockdown of *nlp-40* via RNAi in cholinergic neurons did not yield an increase in survival. However, unexpectedly, we find animals expressing *nlp-40* RNAi in cholinergic neurons, compared to control animals, had increased survival to anoxia (Figure 3B, *unc-17p* empty 0.57 +/-0.06 versus *unc-17p::nlp-40* RNAi 0.77 +/-0.05, t()=3.993, p=0.004, paired t-test). Interestingly, animals that expressed *nlp-40* knockdown in cholinergic neurons phenocopied animals with pan-neuronal knockdown of *nlp-40.* We considered the possibility that *nlp-40* may function broadly in the nervous system to regulate anoxia. If this is the case, knockdown of *nlp-40* in any population of neurons will confer protection against anoxia. We tested knockdown of *nlp-40* in glutamatergic neurons because many neurons that respond to changes in oxygen concentration receive and use glutamate [26]. Additionally, hypoxia alters glutamatergic signaling and glutamate-regulated locomotor behaviors [47]. We reasoned that if *nlp-40* functions in glutamatergic neurons to regulate anoxia, then loss of *nlp-40* in these populations of neurons should similarly increase survival to anoxia. To determine if *nlp-40* functions broadly in neurons, with overlapping function in glutamatergic neurons, we performed RNAi-mediated knockdown of *nlp-40* in glutamatergic neurons using the strain XE1582 (*wpSi11 II; eri-1(mg366)IV;rde-1(ne219)V;lin-15(n744)).* However, we find no statistical difference between the animals raised on empty vector bacteria compared to those raised on RNAi bacteria to induce knockdown of *nlp-40* in glutamatergic neurons (Figure 3B, *eat-4p::* empty 0.60 +/-0.03 versus *eat-4p:: nlp-40*RNAi 0.56 +/-0.02, t(5)=1.275, p=0.25, paired t test). Together, these findings suggest that NLP-40 functions in neurons, in part, within cholinergic neurons to regulate anoxia.

**Figure 3:**
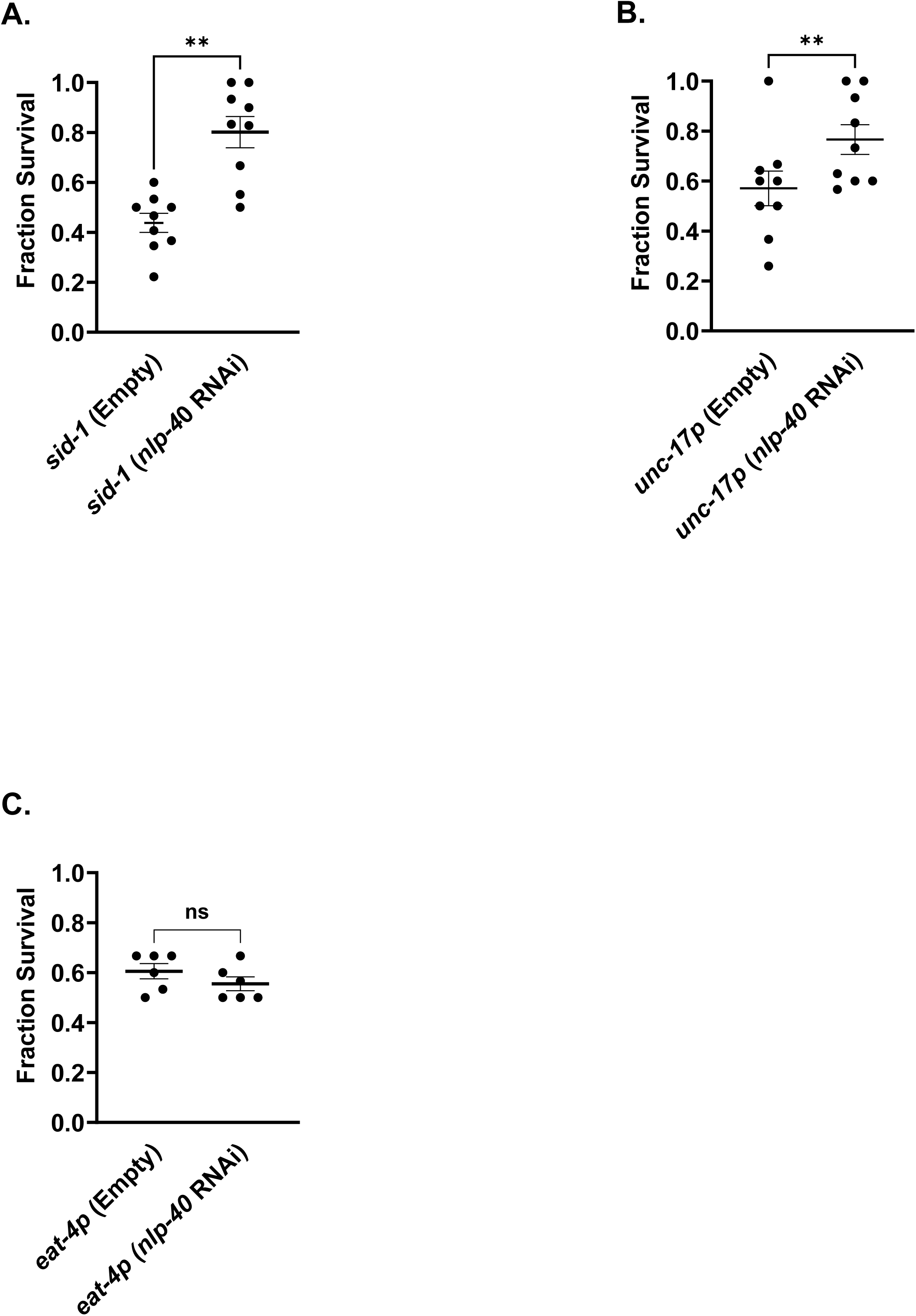
Neuronal Knockdown of NLP-40 Increases Survival to Anoxia. **A)** TU3311 (*unc-119p::sid-1*), a sensitized strain that allows for knockdown of target genes in all neurons or cholinergic neurons, respectively, was fed empty RNAi vector or *nlp-40* RNAi-expressing bacteria for at least 3 generations. L4-stage animals were asphyxiated for 54 hours. Mean fraction of survival is reported for animals that regained movement and/ or feeding after 24 hours of re-oxygenation. 3 independent trials, n= 90 animals per genotype. Error bars represent the standard error of the mean. Statistical significance was analyzed using paired t-tests between empty RNAi and *nlp-40* RNAi for each genotype. * p value= 0.001. **B)** XE1581 (*unc-17p::rde-1*), a sensitized strain that allows for knockdown of target genes in cholinergic neurons, was tested as in panel A. Error bars are the standard error of the mean. Statistical significance was assessed using paired t-tests between empty RNAi and unc-17::*nlp-40* RNAi for each genotype. * p value=0.004 **C)** XE1582 (*eat-4p::rde-1*), L4-stage animals were tested and scored as in panel A. 2 independent trials, =60 animals per genotype. Error bars represent the standard error of the mean. Statistical significance was assessed using paired t-tests between empty RNAi and eat-4p::*nlp-40* RNAi for each genotype.

### NLP-40 and AEX-2 Regulate the Behavioral Response To Anoxia

Animals undergo behavioral changes as part of the homeostatic response to hypoxia [17–19, 26, 48, 49]. In response to low or high oxygen conditions, *C. elegans* increase locomotor speed accompanied by an increase in the frequency of reversals or turning behaviors [17, 25, 26, 50]. In *C. elegans*, neuropeptides can mediate behavioral states by shifting strategies from forward locomotion to reversal turns in response to changes in environmental, chemical, physical, or biological conditions [48, 51, 52]. We asked if NLP-40 or cognate receptor AEX-2 regulated the behavioral response to anoxia. To test whether loss of either *nlp-40* or *aex-2* function altered the behavioral response to anoxia, we scored body bends as a proxy for basal activity, as well as the frequency of reversal turns in L4 animals under normoxia with ample oxygen and under anoxic conditions. Under normoxia, in standard laboratory conditions, we found that L4 *nlp-40* or *aex-2* loss of function mutant animals had reduced body bends compared to stage and age matched wild type controls (Figure 4A, N2 12.8 +/-0.39 versus *nlp-40(tm4085)* 7.3+/-0.3, N2 versus *aex-2(sa3)* 4.3 +/-0.03, ANOVA, F () = 282.4, p value=0.0001). We hypothesized that decreases in either *nlp-40* or *aex-2* function may confer protection against anoxia by reducing activity, either by changing body bends, reversal frequency, or both during anoxia.

**Figure 4:**
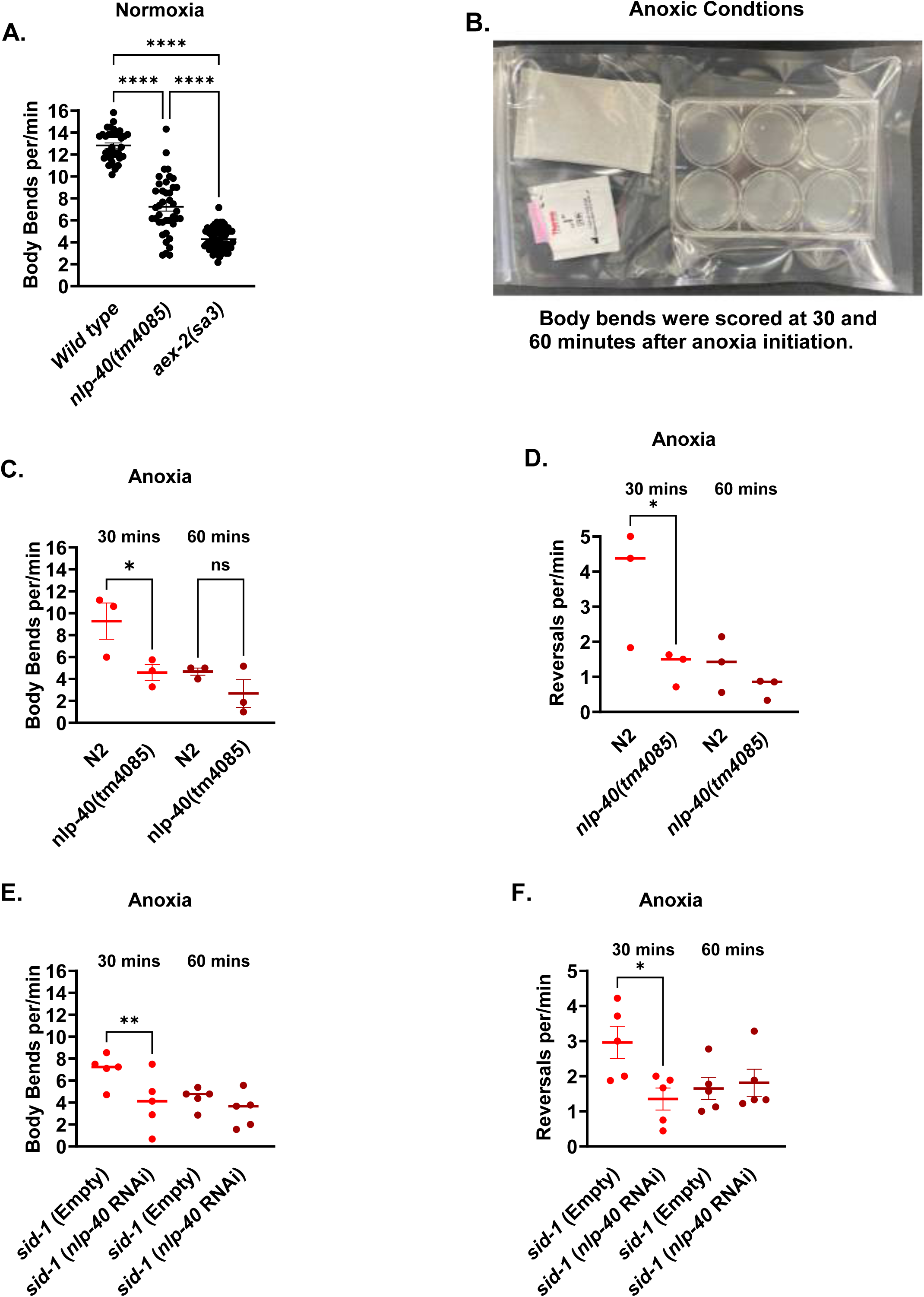
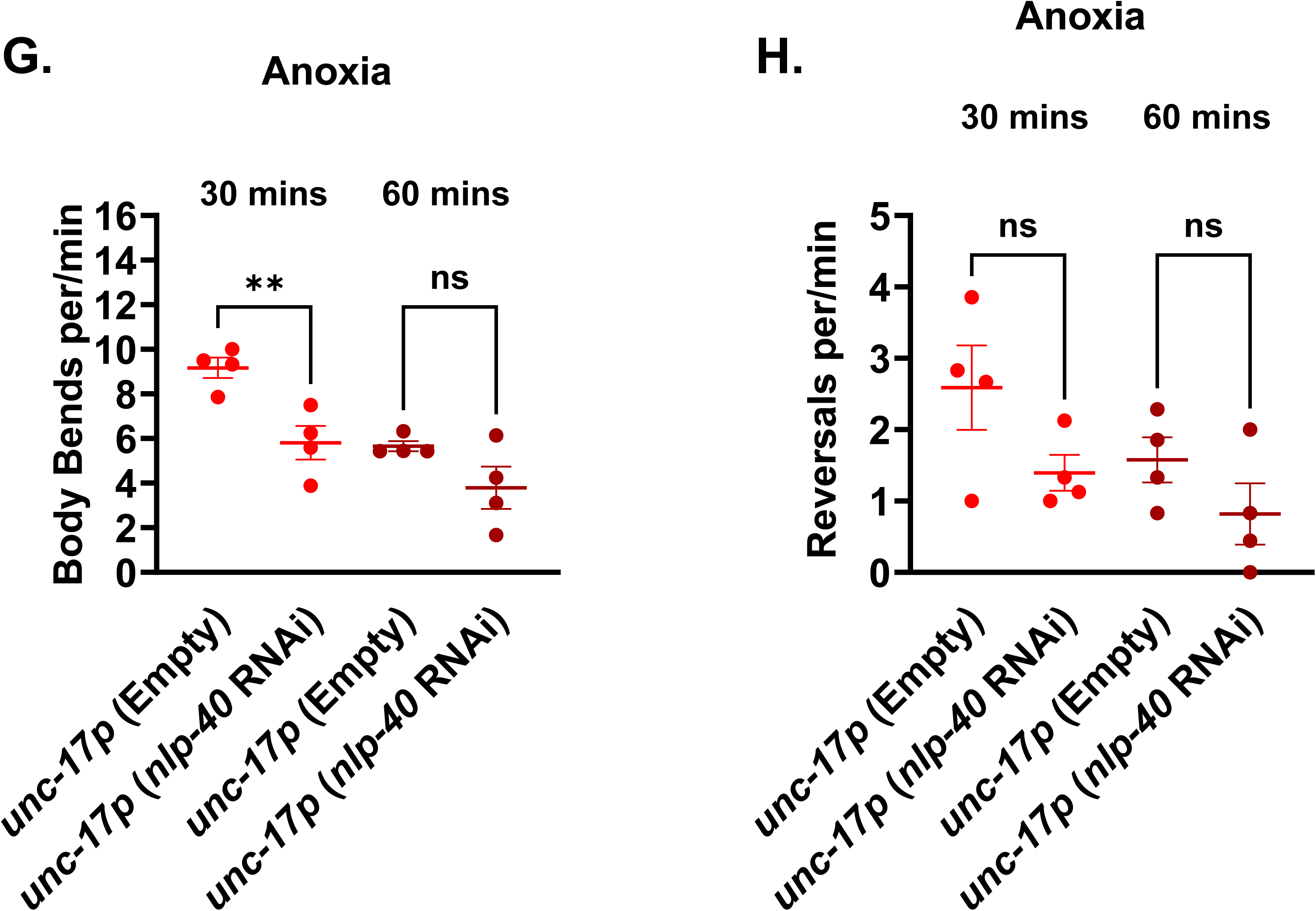
Perturbation in either *nlp-40* or *aex-2* gene function alters the body bend frequency during anoxia. **A) Loss of either *nlp-40* or *aex-2* leads to reduced activity in normoxia in L4-stage animals.** Early L4-stage animals of each strain were selected onto an NGM plate seeded with *OP50 E. coli* the night before testing. Animals were allowed to recover for 3 minutes at laboratory room air conditions, after which the number of body bends was scored continuously for 3 minutes at temperatures between 20 and 22 degrees Celsius. The number of body bends for each strain was then calculated for one minute. 10 animals for each genotype were tested; results are shown for five independent trials (n=50). Statistical significance was assessed by one-way ANOVA with Bonferroni correction, p value < 0.0001. **B) Experimental paradigm for assaying body bends in anoxia.** L4-stage animals are picked to the six-well assay plates. Three animals of the same genotype are placed into each well. Stage- and age-matched controls are tested with experimental strains. Animals are allowed to acclimatize and placed in AnaeroGen anaerobic bag environments for 10 minutes, prior to the start of assays and initiation of anoxia. Body bends were subsequently scored 30 minutes and 60 minutes after the start of anoxia. **C) Loss of neuropeptide *nlp-40* Gene Function Reduced Body Bends During the Beginning Minutes of Anoxia.** Body bends of *nlp-40(tm4085)* L4 stage animals at 30 minutes and 60 minutes after the start of anoxia.*nlp-40(tm4085)* animals compared to wild-type stage- and age-matched controls. Average body bends per minute is reported for 3 independent trials, 27 animals per genotype. Statistical significance was assessed by two-way ANOVA, with a Sidak post hoc correction test, p value 0.02. **D)** Loss of *nlp-40* Decreases Reversal Frequency During Anoxia. Reversal frequency of *nlp-40(tm4085)* L4 stage animals at 30 minutes and 60 minutes after the start of anoxia.*nlp-40(tm4085)* animals compared to wild-type stage- and age- matched controls. The average reversal frequency is reported for 3 independent trials. Statistical significance was assessed by two-way ANOVA, with a Sidak post hoc correction test, p value 0.0005. **E)** *nlp-40* functions in neurons to perturb the number of body bends during anoxia. L4 stage TU3311 (*unc-119p::sid-1*) fed *nlp-40* RNAi expressing bacteria or an empty RNAi vector for at least 3 generations, as in panel A. The average number of body bends per minute shown for 4 independent trials, n= 36 animals per genotype. Statistical significance assessed by two-way ANOVA, with a Sidak correction, p value 0.02. **F)** Knockdown of *nlp-40* in Neurons is Sufficient to Perturb Reversal Frequency. Animals and experimental setup as in panel E. L4 stage TU3311, (*unc-119p::sid-1*) animals tested for reversal frequency at 30 minutes and 60 minutes after initiation of anoxia. The average number of body bends per minute shown for 4 independent trials, n= 36 animals per genotype. Statistical significance assessed by two-way ANOVA, with a Sidak correction, p value 0.02. **G) Knockdown of *nlp-40* in Cholinergic Neurons Reduces Body Bend Activity During Anoxia.** Animals and experimental setup as in panel E. L4 stage XE1581 (*unc- 17p*::empty) fed *nlp-40* RNAi expressing bacteria or an empty RNAi vector for at least 3 generations, as in panel A. The average number of body bends per minute shown for 4 independent trials, n= 36 animals per genotype. Statistical significance assessed by two- way ANOVA, with a Sidak correction, p value 0.0001. **H) Diminished *nlp-40* Expression in Cholinergic Neurons Does Not Alter Reversal Frequency**. Animals and experimental setup as in panel F. L4 stage XE1581, (*unc- 17p*::empty) fed *nlp-40* RNAi tested for reversal frequency at 30 minutes and 60 minutes after initiation of anoxia. The average number of body bends per minute shown for 4 independent trials, n= 36 animals per genotype. Statistical significance assessed by two- way ANOVA, with a Sidak correction, p value 0.09.

Behavior can shift quickly with changes in Oconcentrations [53]. As a result, we evaluated body bends and reversal frequency in *nlp-40* and *aex-2* loss-of-function animals at 30 minutes and 60 minutes after the start of anoxia (Figure 4B). We found that by 90 minutes, all animals were arrested, with cessation of movement (data not shown). To determine if loss of *nlp-40* function altered the behavioral response to anoxia, we tested *nlp-40(tm4085)(lf)* animals in our assay and found that these animals had decreased body bends at 30 minutes compared to wild type stage and age matched controls (Figure 4C, N2 9.2 +/-1.6 versus *nlp-40(tm4085)* 4.6 +/- 0.71, Two-way ANOVA, F()=5.657, p value 0.02). Loss in *nlp-40* also decreased reversal turning frequency compared to wildtype controls (Figure 4D, N2 3.7 +/- 0.96 versus *nlp-40(tm4085)* 1.2 +/- 0.28, Two-way ANOVA, F()=12.20, p-value 0.02). These results suggest NLP-40 regulates the behavioral response to anoxia.

Our results, in combination with others, suggest NLP-40 functions in neurons to regulate response to anoxia [1]. Therefore, we tested whether neuronal loss of *nlp-40* would disrupt body bends and reversal frequency during anoxia, as we observed in *nlp- 40(lf)* animals. We tested TU3311 animals that were fed either *nlp-40* RNAi to reduce *nlp- 40* expression throughout the entire nervous system, or empty vector bacteria as a control. If *nlp-40* functions in the nervous system to mediate body bend activity during anoxia, then we would expect neuronal knockdown of *nlp-40* to disrupt body bend and reversal frequency. Consistent with our logic, we find neuronal knockdown of *nlp-40* led to reduced body bends (Figure 4E, *sid-1* empty 7.0 +/-0.63 versus *sid-1 nlp-40* RNAi 4.0 +/-1.13, Two-way ANOVA, F()=5.543, p value 0.002) and reversal frequency at 30 minutes, compared to control animals fed empty vector (Figure 4F, *sid-1* empty 3.0 +/- 0.46 versus sid-1 *nlp-40* RNAi 1.4 +/-0.38, Two-way ANOVA, F()=7.871 p value 0.0003). Our data suggest *nlp-40(lf)* mutant animals exhibit partial protection from anoxia, in part through reduced activity in the beginning minutes of anoxia. Furthermore, NLP-40 signals from neurons to regulate behavioral responses during anoxia. Next, we tested if inhibiting *nlp-40* expression in cholinergic neurons, would similarly reduce body bend and reversal frequency as observed in animals with pan-neuronal knockdown of *nlp-40*. We found that inhibiting *nlp-40* expression in cholinergic neurons reduced body bend frequency at 30 minutes compared to controls (Figure 4G, *unc-17p* empty 9 +/-0.46 versus *unc-17 nlp-40* RNAi 5.8 +/- 0.75, Two-way ANOVA, F()= 11.50, p value 0.001), but did not perturb reversal frequency (Figure 4H, *unc-17p* empty 2.6 +/- 0.59 versus *unc- 17 nlp-40* RNAi 1.4 +/-0.25, Two-way ANOVA, F()=2.398, p value 0.09). These results suggest that NLP-40 likely functions in cholinergic neurons to regulate the behavioral responses to anoxia.

To further evaluate *nlp-40* in the behavioral response to anoxia, we tested its putative receptor *aex-2*. If *nlp-40* requires aex-2 to regulate the behavioral response to anoxia, loss of *aex-2* should mimic the behavioral responses (decreased body bends and reversal frequency) observed in *nlp-40(lf)* mutant animals. We found *aex-2(sa3)* mutant animals had decreased body bends by 30 minutes compared to wild type controls (Figure 5A, N2 8.4 +/-0.64 versus *aex-2(sa3)* 5.6 +/-0.63, Two-way ANOVA, F()=10.67, p value 0.0002). However, we found no statistical difference in reversal frequency in *aex- 2(sa3)* animals compared to controls (Figure 5B, N2 2.1 +/-0.23 versus *aex-2(sa3)* 1.9 +/- 0.13, Two-way ANOVA, F()=6.159, p-value 0.991). These results suggest that loss of *aex-2* perturbs body bends as early as 30 minutes of anoxia, without changing reversal frequency.

**Figure 5:**
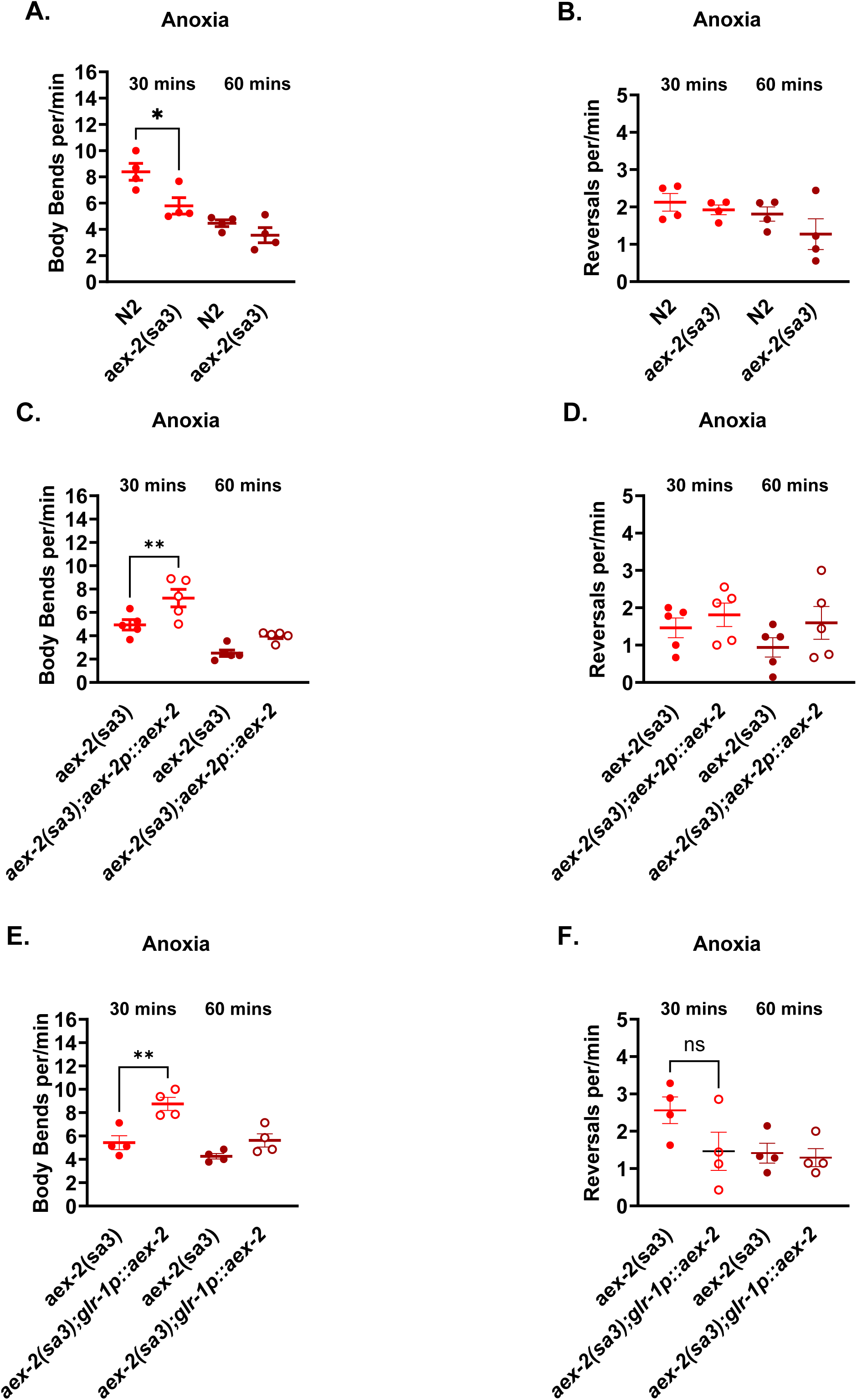
AEX*-2* Function in Neurons to Mediate Body Bend Activity During Anoxia. **A)** Loss in *aex-2* is Sufficient to Reduce the Number of Body Bends During Anoxia. Body bends for *aex-2(lf)* mutant animals were scored at 30 minutes and 60 minutes after the start of anoxia. Stage- and age-matched wild-type animals were tested in parallel. Results shown for an average of 5 independent trials, 45 animals per genotype. Statistical significance was assessed by two-way ANOVA, with a Sidak correction, p value 0.01. **B)** Reduced *aex-2* Function Does Not Perturb Reversal Frequency During Anoxia. Experimental setup, analysis, and presentation as in panel A, but scored for reversal frequency 30 and 60 minutes after initiation of anoxia. Statistical significance assessed as in panel A; no significance. **C) Restoring *aex-2* Behind Its Native Promoter is Sufficient to Rescue the Number of Body Bends During Anoxia.** L4 stage *aex-2(lf)* animals expressing rescue copy of aex-2cDNA behind the native *aex-2* promoter scored for body bends at 30 minutes and 60 minutes after the start of anoxia. Stage-matched *aex-2(sa3lf)* mutant animals tested side by side as controls. Results shown for the average of 5 independent trials, 45 animals per genotype. Statistical significance assessed as in panel A, p value 0.005. **D)** Restoring *aex-2* Function Does Not Rescue Reversal Frequency During Anoxia. Experimental setup, analysis, and presentation as in panel C, but scored for reversal frequency 30 and 60 minutes after initiation of anoxia. Results shown for the average of 5 independent trials, 45 animals per genotype. Statistical significance assessed as in panel A. **E) *aex-2* Gene Function in GLR-1 Neurons Restores the Behavioral Response During Anoxia.** L4-stage *aex-2(lf)* animals expressing rescue copy of aex-2cDNA behind the *glr-1* promoter and *aex-2(lf)* were tested and scored as described in panel D. The average number of body bends per minute is shown for 4 independent trials, 36 animals per genotype. Statistical significance assessed as in panel A, p value 0.007. **F)** Expressing *aex-2* in GLR-1 Neurons Does Not Rescue Reversal Frequency. Experimental setup, analysis, and presentation as in panel E, but scored for reversal frequency 30 and 60 minutes after initiation of anoxia. The average number of body bends per minute is shown for 4 independent trials, 36 animals per genotype. Statistical significance assessed as in panel A.

We already observed that restoring *aex-2* function behind its native *aex-2* promoter was sufficient to restore susceptibility to anoxia (Figure 2A). We reasoned that restoring *aex-2* function behind the native *aex-2* promoter (*aex-2p::aex-2cDNA*) would also restore the number of body bends during anoxia. As we expected, restoring *aex-2* function behind its native promoter, in all tissues where *aex-2* is expressed, rescued body bends frequency, at 30 minutes (Figure 5C, *aex-2(sa3),* 5 +/-0.43 versus *aex-2(sa3);aex- 2p::aex-2,* 7.2 +/- 0.75, Two-way ANOVA, F()=12.80 p value 0.005), but did not increase reversal frequency compared to *aex-2(sa3)* mutant controls (Figure 5D, *aex- 2(sa3),* 1.5 +/- 0.26 versus *aex-2(sa3);aex-2p::aex-2,*1.8 +/- 0.31, Two-way ANOVA, F()=3.482, p value 0.49). Importantly, in these assays, rescue animals had decreased survival compared to *aex-2(sa3)* mutant animals (data not shown), suggesting AEX-2 is necessary to regulate body bend locomotor activity. Next, we wanted to interrogate the cells where AEX-2 functions to regulate body bend activity during anoxia. Because *glr-1* is expressed in many motor neurons and interneurons, including those in neural circuits that control locomotion [54–57], and *glr-1* interneurons are implicated in *C. elegans* Obehavioral responses [58], we prioritized testing *aex-2* rescue in these neuron populations. To determine if restored *aex-2* function in *glr-1* neurons was sufficient to rescue body bend activity during anoxia, we tested *glr-1p::aex-2cDNA* animals. We found that restoring *aex-2* function in *glr-1* expressing neurons was sufficient to rescue the number of body bends in the first 30 minutes of anoxia (Figure 5E; *aex-2(sa3),* 5.4 +/- 0.59 versus *aex-2(sa3);glr-1p::aex-2,* 8.6 +/- 0.55, Two-way ANOVA, F(5,15)=8.623 p value 0.007), but again we saw no significant change in the frequency of reversal frequency in the *glr-1p::aex-2* rescue compared to *aex-2(sa3)* loss of function animals (Figure 5F; *aex-2(sa3),* 2.6 +/-0.36 versus *aex-2(sa3);glr-1p::aex-2,* 1.5 +/- 0.51, Two-way ANOVA, F(5,15)=4.586 p value 0.08). Together, our results support a model in which *aex-2* partly functions in *glr-1* neurons to regulate anoxia, and restoring *aex-2* function in these neuron populations is sufficient to mediate body bend activity during anoxia.

## Discussion

In response to changes in oxygen availability, animals modify their movement strategies to escape and find more suitable conditions. *C. elegans* respond to changes in oxygen concentration with increased speed in locomotion and increased frequency of reversals[17, 18, 22, 26]. In this study, we investigated a role for neuropeptide *nlp-40* and its putative receptor *aex-2* in regulating these behaviors in response to anoxia. Consistent with previous findings, we found that loss of either *nlp-40* or its cognate receptor *aex-2* increased survival to 48 hours of anoxia. We show that loss of either *nlp-40* or its receptor, *aex-2*, decreases escape behaviors (body bends), which protect against anoxia. We found that NLP-40 signals from cholinergic neurons and AEX-2 signals from interneurons or motor neurons render animals susceptible to anoxic insult.

We hypothesized that deficiency in either *nlp-40* or *aex-2* conferred survival by altering the behavioral responses to anoxia. We tested this hypothesis by exposing *nlp- 40(lf)* or *aex-2(lf)* mutant animals to anoxia and counted body bends, as well as reversal frequency, as a measure of activity. We found that *nlp-40(lf) and aex-2(lf)* animals had a reduced number of body bends compared to wild-type animals 30 minutes after the start of anoxia. *nlp-40(lf)* animals also had reduced reversal frequency. However, unlike *nlp- 40(lf)* animals, we did not find statistical differences in reversal frequency in *aex-2(lf)* mutant animals compared to controls. One interpretation is that AEX-2 does not function in neurons and circuits to regulate reversals. Depending on environmental or cellular conditions, distinct neuropeptides and receptor signaling pathways determine specific behavioral responses [31]. We consider a similar mechanism here: NLP-40 may regulate body bend activity through AEX-2 and reversal frequency through AEX-2-independent mechanisms. We cannot exclude the possibility that mutations in *aex-2* alter reversal frequency and our experimental setup may limit our ability to capture subtle changes in behavior. Nevertheless, we consistently detect changes in body bend frequency and overall activity in *nlp-40* and *aex-2* mutant animals. Combined, our results support a model in which NLP-40 functions partially through AEX-2 to regulate aspects of the behavioral response, specifically body bend activity, to anoxia.

Several groups have shown a neuronal role in mediating the behavioral response to changes in oxygen [52, 59]. We and others have found that inhibiting *nlp-40* expression broadly in all neurons increases survival to anoxia [1]. We find that NLP-40 signals are specific to neuron populations; diminished *nlp-40* expression in glutamatergic neurons does not increase survival to anoxia, demonstrating cell specificity in response. Doshi et al. found that inhibiting *nlp-40* gene expression from cholinergic neurons did not increase survival to anoxia. However, unexpectedly, we show that loss of *nlp-40* gene expression in cholinergic neurons confers an advantage and increases survival to anoxia. Our experimental setup and design are similar to the previous study, and the source of discrepancy is unclear; however, we speculate that subtle differences in laboratory conditions may contribute in part to differences between our results and those published in Doshi et al., 2019. dsRNA against a targeted gene to reduce gene expression produces variable knockdown effects [38, 60–62]. Under standard laboratory conditions, we achieved a more robust reduction in gene expression.

Regardless, our results support a role for NLP-40 function in cholinergic neurons. We used bioinformatic platforms to identify neurons where *nlp-40* is expressed to uncover possible candidates (Figure 6). Among the candidate neurons are the ASK neurons; however, Doshi et al. showed that inhibiting *nlp-40* expression in these neurons did not confer protection against anoxia. We consider other possible neurons based on the following criteria: (i) neurons that either use or release acetylcholine neurotransmitter input and (ii) inhibition of the candidate neuron(s) led to similar behavioral defects that were observed in our *nlp-40* mutants. Based on these criteria, RIM neurons emerge as candidates for where NLP-40 may function to regulate anoxia. Behavioral studies show laser ablation of RIM leads to either increased or decreased reversal frequency [51, 63], suggesting RIM has both activating and inhibitory roles within the neural circuit and provides support for exploring RIM as a neuron of interest. Another candidate of interest include AIB command interneurons. AIB is embedded in a neural circuit that drives reversals in response to noxious or aversive stimuli [64, 65]. Alternatively, it is possible that NLP-40 functions more broadly, in cholinergic neurons, in neural circuits that regulate locomotion.

**Figure 6:**
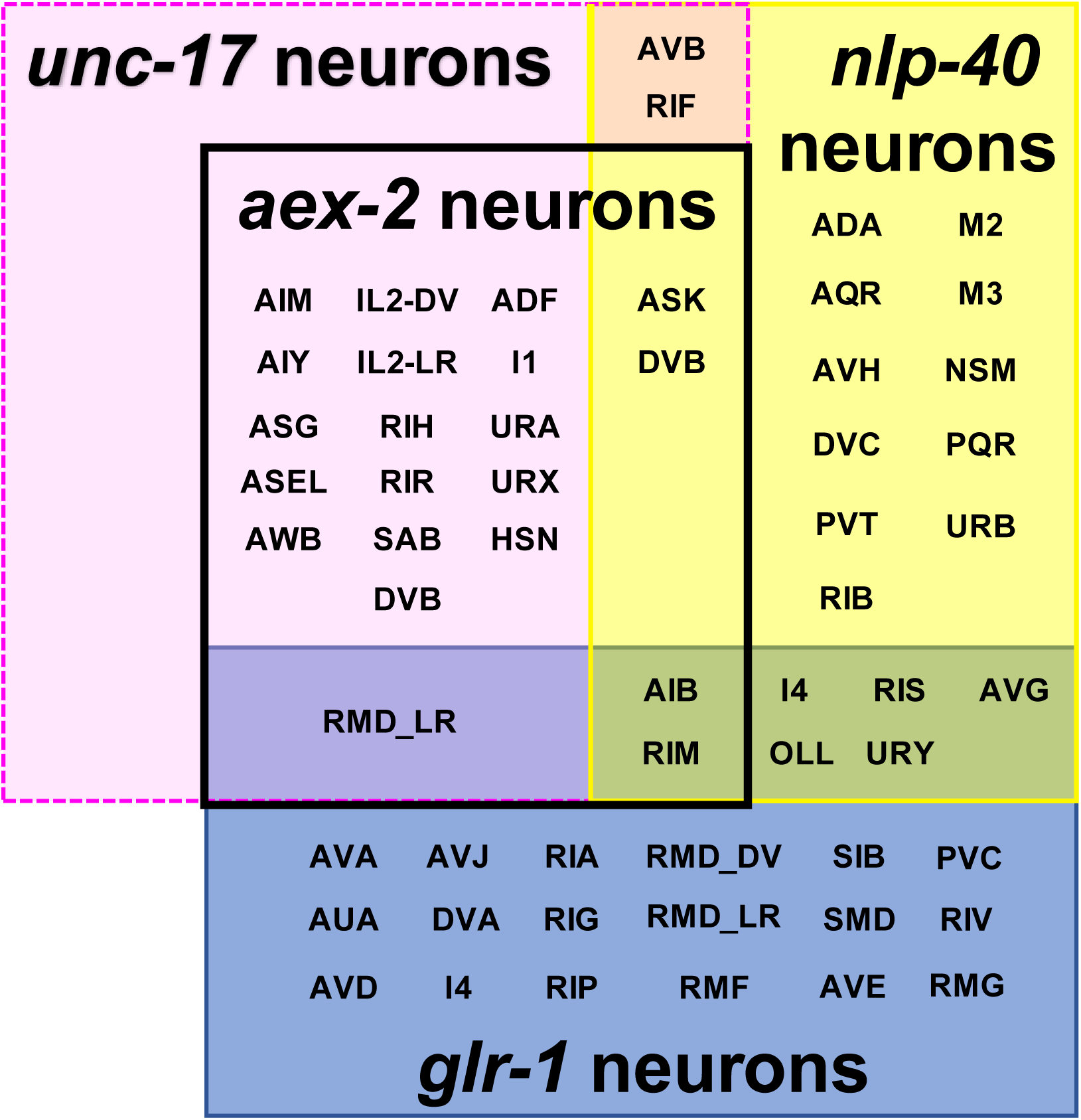
NLP-40 and AEX-2 Likely Function in GLR-1 Neurons To Regulate Behavioral Responses to Anoxia. *Caenorhabditis elegans* Neuronal Gene Expression Map and Network (CeNGEN) was used to identify all neurons containing RNA transcripts for either *nlp-40*, *aex-2*, *glr-1* promoter, or *unc-17* promoter. For *nlp-40*, approximately 12 neurons with overlapping expression of either *glr-1*, *unc-17, or aex-2*. Figure includes all *nlp-40 and glr-1* neurons, but not all *aex-2* or cholinergic/*unc-17* neurons. Among candidate neurons, GLR-1 interneurons RIM and AVB command interneurons contain *nlp-40* transcripts. For *aex-2*, we identified RIM neurons that express *glr-1* and *aex-2*, and RMD_LR neuron pairs that express *aex-2*, *glr-1*, and *unc-17* as strong candidate neurons to test for cellular sites of action for AEX-2 in regulating responses to anoxia. Neurons that also express the GABAergic promoter *unc-47* were deprioritized from consideration, as they did not restore vulnerability to anoxia in the *aex-2(lf)* mutant animals, and likely not where AEX-2 functions to regulate the behavioral responses to anoxia.

We tested tissue specificity for *aex-2* by restoring gene function in neuronal and non-neuronal cells using extrachromosomal array strains that express *aex-2cDNA* behind tissue-specific promoters. We find that restoring *aex-2* function in both cholinergic and a subset of motor and interneurons increases vulnerability to anoxia. One important consideration is that *unc-17* and *glr-1* promoters, which drive expression broadly in acetylcholine-releasing neurons and a subset of motor and interneurons, respectively, overlap in expression in 14 neurons [45], and may account for restored vulnerability to anoxia observed in both strains. We found that restoring *aex-2* function in *glr-1* neurons rescued body bend activity in *aex-2(lf)* mutant animals. RNA sequencing analyses and bioinformatic platforms indicate that approximately thirty-three neurons express *aex-2* RNA transcripts (Supplemental Table 1) [45, 66]; however, few neurons express both *glr- 1* and *aex-2*. Furthermore, only one neuron pair, RMD_LR, expresses *unc-17*, *aex-2*, and *glr-1* and would allow for rescue in both cholinergic and *glr-1* neuron populations (Figure 6). RMD is embedded in a neural circuit that regulates spontaneous locomotion for exploratory foraging behaviors, and inhibition of RMD leads to less movement [54], making RMD a strong candidate to test as a possible cell for where *aex-2* functions for anoxia. Moreover, RIM forms chemical synaptic connections onto RMD, in part through neuropeptides [67], and bolsters support for their involvement in regulating escape behavioral responses to anoxia [68, 69]. It is possible that AEX-2 functions more broadly in cholinergic motor and GLR-1 interneuron locomotor circuits. Future studies will further interrogate the neural mechanisms by which neuropeptide *nlp-40* and *aex-2* regulate the behavioral responses to changes in hypoxic states.

Finally, we consider what might confer protection and increase survival to anoxia. We show that loss of either *nlp-40* or its receptor, *aex-2,* decreased escape behaviors and body bends, and this provided protection and increased survival to anoxia. We found that NLP-40 signals from cholinergic neurons and AEX-2 signals from interneurons or motor neurons render animals susceptible to anoxic insult. How does reduced NLP-40 and AEX- 2 signaling confer resistance to anoxia? In the wild, increased locomotor speed, reversals, and turning behaviors are protective and allow animals to escape to find more favorable conditions. However, in standard laboratory settings, *C. elegans* have limited options for escape when confronted with unfavorable conditions, and these same behaviors are deleterious and make animals more vulnerable to anoxia. The simplest model is that NLP- 40 and AEX-2 release, likely from interneurons or motor neurons, increases the activity of neurons that drive escape behaviors that would be beneficial in the wild, but in confined conditions, are harmful. One interpretation is that changes in behavioral activity within a subset of neurons lead to inherent sensitivity or resistance to changes in oxygen concentrations. However, this seems unlikely, and we favor an interpretation where reduced NLP-40 and AEX-2 release also leads to changes in gene expression in downstream target cells that may render animals more resistant and confer protection. We cannot exclude the possibility that less neuropeptide release from neurons affects metabolism and may also contribute to protection against anoxia. Understanding the molecular changes underlying changes in behavior and how neuropeptide signaling affects the response will be of great interest and will expand our knowledge how animals adapt to changes in low oxygen environments.

## Conclusions

The molecular and neural mechanisms underlying the responses to low oxygen conditions have not been fully described. We and others find that loss of *C. elegans* neuropeptide *nlp-40* or *aex-2* G-protein-coupled receptor confers protection and increased survival to 48 hours of anoxia. Here we report that NLP-40 and AEX-2 function in neurons, partially in cholinergic and GLR-1 interneurons or motor neurons, to mediate the adaptive behavioral responses to anoxia. Our results shed light on mechanisms that regulate adaptive behavioral responses to oxygen-deprived conditions and highlight a novel role for NLP-40 and AEX-2 in regulating aspects of this homeostatic response.

## Data Availability

All data that contributed to the findings, including the number of animals scored and the number of independent replicates, are included in the supplement. *C. elegans* strains or plasmids generated or used in this study are available upon request.

## Acknowledgements

We thank Michael Nonet and Derek Sieburth for generously sharing several strains, plasmids, and reagents used in our experimental paradigm. Some strains were provided by the CGC, which is funded by the NIH Office of Research Infrastructure Programs (P40 OD010440) and the Japanese National Bioresource Project. We also thank Anne C. Hart at Brown University and the labs in the Yale University Worm Group, for helpful discussions. We also thank members of the Bennett lab at Trinity College: Bettina Evans, Rothswell Lanting, and Emmett McGranaghan for feedback and discussions.

## Author Contributions

H.L.B. conceptualized and designed the experiments and wrote the manuscript. G.Z.W. and H.L.B. conducted experiments and prepared figures. G.Z.W. and H.L.B. analyzed all results. All authors proofread the final manuscript.

## Notes

### Competing Interest Statement

The authors have declared no competing interest.

